# A tissue dissociation method optimized for ATAC-seq and CUT&RUN in *Drosophila* pupal tissues

**DOI:** 10.1101/2022.08.15.504052

**Authors:** Elli M. Buchert, Elizabeth A. Fogarty, Christopher M. Uyehara, Daniel J. McKay, Laura A. Buttitta

**Affiliations:** Molecular, Cellular and Developmental Biology, University of Michigan, Ann Arbor 48109; Dept. of Biology, Dept. of Genetics, Integrative Program for Biological and Genome Sciences, University of North Carolina, Chapel Hill, Chapel Hill, NC 27955; Curriculum in Genetics and Molecular Biology, University of North Carolina, Chapel Hill, Chapel Hill, NC 27955

## Abstract

Chromatin accessibility, histone modifications and transcription factor binding are highly dynamic during *Drosophila* metamorphosis and drive global changes in gene expression as larval tissues differentiate into adult structures. Unfortunately, the presence of pupa cuticle on many *Drosophila* tissues during metamorphosis prevents enzyme access to cells and has limited the use of enzymatic *in situ* methods for assessing chromatin accessibility and histone modifications. Here, we present a dissociation method for cuticle-bound pupal tissues that is optimized for use with ATAC-Seq and CUT&RUN to interrogate chromatin accessibility and histone modifications. We show this method provides comparable chromatin accessibility data to the non-enzymatic approach FAIRE-seq, with only a fraction of the amount of input tissue required. This approach is also compatible with CUT&RUN, which allows genome-wide mapping of histone modifications with less than 1/10^th^ of the tissue input required for more conventional approaches such as Chromatin Immunoprecipitation Sequencing (ChIP-seq). Our protocol makes it possible to use newer, more sensitive enzymatic *in situ* approaches to interrogate gene regulatory networks during *Drosophila* metamorphosis.

## Introduction

The study of global gene regulatory networks during development requires observational data describing changes in chromatin accessibility, histone modifications and transcription factor binding over time, in a manner coordinated with cellular differentiation. Molecular studies of developmental gene regulatory networks *in vivo* have often involved pooling of embryos, dissection of tissues and/or sorting of specific cell types to isolate nuclei or chromatin, followed by nuclease assays or Chromatin ImmunoPrecipitation (ChIP) (Wilczynski and Furlong, 2010). Newer enzymatic methods to examine the molecular underpinnings of gene regulation such as Assay for Transposase-Accessible Chromatin with high-throughput sequencing (ATAC-seq) and Cleavage Under Targets and Release Using Nuclease (CUT&RUN) have made it possible to assay chromatin accessibility and histone modifications or transcription factor binding with a fraction of the input required for more traditional assays (Ahmad and Spens, 2019; Davie et al., 2015), making them ideal tools for experiments that require tissue dissection or cell sorting (Uyehara and McKay, 2019). But these assays can be challenging in tissue types where the cells are heavily embedded in the extracellular matrix, or in the case of insects, enclosed within a tough cuticle, which limits nuclear isolation by homogenization and enzyme access to cells.

We previously customized a flow cytometry protocol for cuticle-encased tissues such as *Drosophila* pupal wings, to measure DNA content and fluorescent transgene expression (Flegel et al., 2013). Our protocol was designed to dissociate the wing after pupa cuticle deposition, but optimized cell recovery, viability and access to vital DNA dyes for DNA content quantification. We therefore reasoned that with further optimization, our dissociation approach could provide enzyme access to live cells from these tissues to take advantage of newer chromatin profiling techniques toward the study of chromatin dynamics in the fly wing during metamorphosis.

Here, we describe a dissociation method for cuticle-bound pupal tissues such as the *Drosophila* wing, that is optimized for use with subsequent ATAC-Seq or CUT&RUN applications to interrogate chromatin accessibility and histone modifications.

## Methods

### *Drosophila* genotypes and staging

*Drosophila* tissues were dissected from *w^1118^/y,w,hsflp^122^; +; +* female animals for all samples except the CUT&RUN 24h APF 45 and 75 wing samples, which were a mixture of *y,w* males and females. Animals were raised at 25°C on Bloomington Cornmeal media without malt extract (bdsc.indiana.edu/information/recipes/bloomfood.html). Larval samples were dissected from wandering larvae isolated from uncrowded vials. Vials with more than ~100 larvae were diluted into fresh vials to keep larvae uncrowded. Pupa were collected from vials at the White Pre-pupa stage (WPP) as described (Flegel *et al.,* 2013), which was taken as 0h After Puparium Formation (APF) and reared on damp Kimwipes at 25°C to the indicated hours APF.

### Tissue Isolation and Dissociation

We use Collagenase/Dispase from Sigma (cat# 10269638001), which is Ca^2+^ dependent and acts at low temperatures such as 4°C. Collagenase/Dispase is not inhibited by serum, but is compatible with the wash solutions for ATAC-Seq and CUT&RUN and is inhibited by EDTA.

#### Reconstitute Collagenase/Dispase mixture

Collagenase/Dispase comes as 100mg lyophilized powder. To reconstitute, dissolve powder in 1mL dH2O to make 100mg/mL concentrated stock. This can be stored at - 20°C, but we suggest making aliquots of 2mg/mL 2X working stock to avoid repeated freeze/thaw cycles. To make the 2X working stock, dilute the concentrated 100mg/mL stock 1:50 in Wash solution (20 mM HEPES, pH 7.5, 150 mM NaCl, 0.1% BSA). Aliquot 100μL per tube to make multiple aliquots of 2X working stock to avoid repeated freeze thaw cycles.

#### Dissociation

We dissect tissues in glass embryo dishes (Electron Microscopy Sciences) in Wash+ solution (Wash solution supplemented with 0.5 mM Spermidine and Roche complete Protease Inhibitor Tablet 1 tablet per 50 mL). Videos of pupal wing dissections are published (Flegel *et al.,* 2013) and also available upon request. In brief for pupa wing dissection, we use forceps to hold the pupa at the anterior operculum and use microdissection scissors (Fine Science Tools, Vannas Tubingen spring scissors) to perform a posterior cut across the pupa to release fat body and hemolymph. We then perform a second, longitudinal cut on the ventral side of the pupa, extending from the posterior edge to the head region. Forceps can now be used to remove the fileted pupa from the tanned cuticle. The shiny, clear pupa cuticle will now be evident, and we use a glass Pasteur pipet to gently flow dissection solution throughout the opened pupa to remove any remaining fat body. We then pull wings off from the cleaned pupa at the hinge, using sharp forceps (Fine Science Tools, Dumont #5). For some CUT&RUN experiments here (those with 24h APF wings), we also dissected and added 4 *Drosophila virilis* 3^rd^ larval instar wing discs to the dissociation mixture, to use as a spike-in control to verify similar fragment recovery rates across dissociated samples with varying inputs from 20-75 pupal wings.

We use 10 larval wing discs or pupal wings for ATAC-Seq and 20-75 discs or wings for CUT&RUN. The amount of tissue used for CUT&RUN depends on the antibody and must be determined empirically. Keep track of how many tissues you add per sample, particularly when optimizing the protocol and preparing samples for flow cytometry. This will allow for calculations of viable cells released per tissue. Pre-coat a cutoff P200 tip by pipetting up and down in histolyzed fat body from dissected pupa or larva carcasses (to avoid tissue sticking). Pipet 100μl of dissected tissues in Wash+ solution directly into 100μL 2X Collagenase/Dispase solution. Incubate in a thermomixer at 23°C for 30 minutes, shaking at 500 rpm. Vortex for 10 seconds at setting 6 (about 60% of max speed). Incubate another 10 minutes at 23°C, shaking at 500 rpm. Vortex again for 10 seconds at 60% speed. At this point you may see the empty, clear pupa wing cuticle floating at the top of the solution. Proceed to downstream protocols as described for ATAC-seq, CUT&RUN or flow cytometry.

### Flow Cytometry

In pilot experiments used to assess and optimize the dissociation protocol, we measured cell number and viability by flow cytometry. A live cell DNA stain is used to count cells and propidium iodide is used to differentiate live from dead or dying cells. After dissection but before beginning the incubations at 23°C with shaking, add an additional 300 μL of Wash+ solution for a total volume of 500μL. Add 0.5μL of live cell DNA stain (Vybrant DyeCycle Violet DNA Stain, Invitrogen or Hoechst 33342, Sigma) and 1.2 μL of 10mg/mL Propidium Iodide (PI, Sigma). Proceed with shaking at 23°C and vortexing as described above. After the final 10 second vortex, add 500 μL of 1X PBS + 0.1% BSA bringing final sample volume to 1mL. Do not pipet to mix, as cells will stick to the plastic pipet; adding the solution will sufficiently mix the sample. Run sample immediately through a flow cytometer to measure cell number per tissue (gating on diploid 2N and 4N cells stained with live DNA dye) and cell viability (PI positive cells are permeable dead or dying cells, PI negative cells are viable). Our flow cytometry was performed using an Attune NxT at a flow rate of 100μL/min. We consider samples with >95% of PI negative diploid cells to be successful.

### ATAC-Seq

We use the Omni-ATAC Protocol described in (Corces et al., 2017). Dissect and dissociate larval or pupal tissues (10 wings per sample) as described above. After the final 10 second vortex, spin down tissues at 800 x g, for 5 minutes, at 4°C. Remove supernatant and wash in 200 μL Ca^2+^ free 1X PBS. Repeat 4°C spin and remove supernatant. Resuspend cell pellet in 50 μl cold ATAC-RSB (10mM Tris-HCl pH 7.4, 10 mM NaCl, 3mM MgCl_2_) supplemented with 0.1% NP40, 0.1% Tween-20, and 0.01% Digitonin for cell lysis. Pipette up and down 3 times. Incubate on ice 3 minutes. Quench lysis by adding 1 mL cold ATAC-RSB containing 0.1% Tween-20 without NP40 or Digitonin. Invert tube 3 times to mix. Spin down at 800 x g for 10 minutes, 4°C. Discard supernatant and immediately continue to transposition reaction. Note that the pellet can be quite loose at this point. We often remove 1 mL of supernatant, then spin down again for 5 minutes to re-pellet and remove the final 50 μL.

#### Transposition reaction and purification

To make the transposition reaction mix, combine the following per sample: 25 μl 2x TDE1 Buffer (Illumina Cat #20034197), 3.5 μl Tn5 Enzyme (100nM final, Illumina Cat #20034197), 16.5 μl PBS, 0.5 μL 1% Digitonin, 0.5 μL 10% Tween-20, 5 μL water. Resuspend nuclei in 50 μL transposition reaction mix. The transposition reaction is carried out at 37°C for 30 minutes shaking at 1000 rpm. Following transposition reaction, the sample is purified using a Qiagen MinElute kit according to manufacturer’s protocol. Elute transposed DNA in 21 μl Elution Buffer (10 mM Tris buffer pH 8.0). Purified DNA can be stored at −20°C.

### CUT&RUN

Our pupal CUT&RUN protocol is adapted from (Uyehara and McKay, 2019). Dissect tissues (20 wings per sample) in cold CUT&RUN Wash+ buffer, and proceed with Collagenase/Dispase dissociation as described above.

#### Binding dissociated cells to Concanavalin A (ConA) beads

Aliquot 15 μL of Concanavalin A (ConA) beads (Polysciences cat#86057-3) and bind to magnet for 5 minutes, then remove supernatant. Wash beads 2X with 1mL Binding Buffer (20mM HEPES-KOH, pH 7.9, 10mM KCl, 1mM CaCl_2_, 1mM MnCl2), then resuspend in 15 μL Binding Buffer. Transfer dissociated wings to ConA beads and mix with gentle pipetting, then add 1mL DBE (Digitonin Block EDTA: 2mM EDTA, 0.1 % Digitonin in Wash + buffer) and pipette to mix. Incubate for 10 minutes on ice. Bind cells and beads to magnet for 2 minutes. Remove buffer and replace with 100μL DBE supplemented with primary antibody of interest (data presented here: rabbit anti-H3K27Me3 antibody (Cell Signaling, 1:100). Incubate cells plus antibody angled sideways on orbital shaker, set to low, overnight at 4°C. Bind beads to magnet for 2 minutes at 4°C. Remove supernatant and wash 2 X with 500μL DBE, inverting tube back & forth ~10X to mix.

We used two sources of protein A/G-MNase in this study. For larval wings (L3) and 24h pupal samples with 20 wings, we used protein A/G-MNase from Epicypher. For 24h pupal samples with 40 and 75 wings, we used protein A/G-MNase kindly provided by Dr. Steven Henikoff (Meers et al., 2019). For experiments using pA/G Mnase from Epicypher, dilute pA/G Mnase 1:20 into DBE and keep on ice until ready for use. Bind samples to magnet for 2 minutes then remove supernatant and resuspend in 100 μL Protein A/G-MNase solution. Incubate for 10 minutes at room temperature on an orbital shaker, set to low, at an angle. At 4°C, bind samples to magnet for 2 minutes, then remove supernatant. Resuspend beads in 500 μL Wash+ and incubate for 2 minutes. Bind samples to magnet for 2 minutes, then repeat wash once. Remove the supernatant and resuspend samples in 75 μL Digitonin Buffer (Wash + solution with 0.1% Digitonin). Add 75 μL 2x Rxn Buffer (Wash + buffer with 4mM CaCl_2_) to samples. Digest for 2 hours at 4°C on orbital shaker. Add 150 μL 2xSTOP buffer (20 mM HEPES, pH7.5, 200 mM NaCl, 20 mM EDTA, 4 mM EGTA) + 60 μg/ml RNAseA to stop digestion reaction and pipette to mix. For L3 wing samples, a spike-in control was used of *E. Coli* DNA (Epicypher cat# 18-1401) at 5.25 pg per sample. For experiments using pA/G Mnase provided by the Henikoff lab we made the following modifications to the above process; cells were incubated with pA/G Mnase for 1h at 4°C prior to adding Rxn buffer. Digestion was performed in Rxn buffer for 30 min at 4°C.

#### Fragment Release and purification

Incubate samples in thermomixer set to 37°C without shaking for 30 minutes, then bind sample on magnet for 2 minutes. Transfer supernatant to low retention tube labeled “S” (for supernatant). Resuspend the remaining pellet in 150 μL Pellet Buffer (Wash + buffer with an equal volume of 2xSTOP + 10% SDS and 6.89 μL proteinase K 20mg/mL). Add 2 μL 10% SDS and 2.5 μL proteinase K (20mg/mL) to supernatant samples. Mix by briefly vortexing. Incubate supernatant and pellet samples in thermomixer set 50°C without shaking for 2 hours. Place tubes on magnet for 2 minutes, then transfer supernatant to new low retention tube. Fragments were selected using SPRI beads and library preparation was performed using Takara ThruPLEX DNA-seq kits.

#### Sequencing platforms and read depth

Library quality was assessed with an Agilent Tape Station. H3K27me3 CUT&RUN 24h APF 20 wings sample was sequenced using Illumina NextSeq 100 Cycles HO, paired-end 50bp reads, at a target depth of 15 million reads. 45 and 75 wings samples were sequenced on an Illumina NovaSeq SP 100 cycle flow cell for 50 bp paired-end reads, at a target depth of 10 million reads per sample. L3 H3K27me3 CUT&RUN samples were sequenced on an Illumina NovaSeq S4 300 cycle flow cell for 150 bp paired-end reads, at a target depth of 50 million reads per sample. L3 wing disc ATAC-Seq libraries were sequenced on an Illumina NextSeq MO 150 cycle flow cell for 75bp paired end reads, with a target depth of 16 million reads per sample. Pupal wing ATAC-Seq libraries were sequenced on an Illumina NovaSeq SP 100 cycle flow cell for 50bp paired end reads, at a target depth of 90 million reads per sample.

#### Data Analysis

Raw sequencing files were assessed using FastQC (https://www.bioinformatics.babraham.ac.uk/projects/fastqc/), then adaptors and low-quality bases were trimmed using Trimmomatic-0.39 (Bolger et al., 2014) for CUT&RUN libraries, or cutadapt 1.18 (Martin, 2011) for ATAC-Seq libraries. Reads were aligned to DM6 using Bowtie2.4.1 (Langmead and Salzberg, 2012) using --local --very-sensitive parameters and max fragment size set to 1000. PCR duplicates were marked using picard-tools 2.8.1 MarkDuplicates (“Picard Toolkit.” 2019. Broad Institute, GitHub Repository. https://broadinstitute.github.io/picard/; Broad Institute). BAM files were generated using samtools 1.5 (Danecek et al., 2021), and peaks were called using macs2 version 2.1.2 (Zhang et al., 2008). ATAC-Seq fragments spanning less than 120bp (sub-nucleosomal fragments) were used for analysis. Within each analysis (dissociated vs non-dissociated L3 ATAC, ATAC vs FAIRE, etc) library depth was normalized by downsampling larger datasets prior to peak calling. Only peaks that were identified in all replicates for a given timepoint or condition were used in downstream analyses. Bigwig tracks and CUT&RUN heatmaps were generated using DeepTools utilities (Ramirez et al., 2016). ATAC-Seq and FAIRE heatmaps were generated using R package pheatmap version 1.0.12. ATAC-Seq and FAIRE peaks mapping to blacklist regions (Amemiya et al., 2019) and LINE/LTR repeat regions (Karolchik et al., 2004) were excluded from analyses. Read coverage within peaks was calculated using featureCounts from subread version 1.6.0 (Liao et al., 2014). Peaks were assigned to genomic features using R package ChIPpeakAnno (Zhu et al., 2010). Conservation scores for ATAC-Seq and FAIRE peaks were extracted from the UCSC Genome Browser dm6 124-way PhastCons data file (Kent et al., 2002) using a publicly available custom script written by Dr. Ian Donaldson, University of Manchester (https://www.biostars.org/p/16724/#16731).

#### Data Access

Data generated in these studies is available in GEO at GSE211269.

Previously published larval and pupal wing FAIRE data can be accessed from GEO GSE131981.

Previously published L3 wing H3K27Me3 ChIP data was accessed from GEO GSE74080.

## Results & Discussion

### A wing dissociation protocol to release cells from cuticle that minimizes cell death and loss

During *Drosophila* metamorphosis, tissues that make the head, notum, wings, halteres, abdomen and legs are covered by a pupal cuticle secreted by the imaginal epithelium beginning at 6-10 hours after puparium formation (APF) (Reed et al., 1975). To study these tissues after 6h APF, the cuticle is typically removed by hand dissection after fixation, to allow access for antibodies and stains for immunofluorescence on these tissues. We previously attempted to perform ATAC-seq on dissected pupal tissues, but found the pupa cuticle inhibited access to the transposase enzyme. We therefore used a chemical approach with fixation and sonication, FAIRE-seq, to interrogate chromatin accessibility changes in pupal wings (Ma et al., 2019). This chemical approach required at least 40 wings per sample, which involved significant manual dissection and limited the resolution of our timecourse. This prompted us to reinvestigate whether ATAC-seq may work on smaller numbers of unfixed pupal tissues that have been dissociated, to release them from the pupa cuticle.

We started from a well-established protocol for flow cytometry that was developed to study cell cycle changes in developing *Drosophila* wings (de la Cruz and Edgar, 2008) and adapted it for various pupal tissues, from stages 18-44 h APF for cell cycle analysis (Flegel *et al.*, 2013). We tested tissue dissociation with various enzymes including trypsin-based digestion, collagenase, dispase and chitinase, as chitin is a major component of the pupa cuticle (Fristrom et al., 1982). In our hands, dispase alone and several concentrations of chitinase failed to dissociate cells from pupal wings. We found that digestion with a commercially available Collagenase/Dispase mixture (see methods) for about 40 minutes at room temperature with gentle shaking optimized cell recovery from unfixed, hand-dissected tissues and minimized cell death (Fig. 1). Key steps to optimize cell recovery and improve consistency from sample to sample include: 1. pre-coating pipet tips (both glass and plastic) with histolyzed fat body from dissected pupa before using them to transfer dissected tissues in order to avoid sticking and tissue loss, 2. transferring tissues from dissection dishes in a Hepes-based wash buffer with minimal solution carryover, and 3. consistent shaking during dissociation. We prefer to use a temperature and speed-controlled Thermomixer (available from Eppendorf) for this purpose. For performing ATAC-seq after tissue dissociation, we gently pellet the cells and wash once with Ca^2+^ free 1X PBS to remove enzymes. For CUT&RUN, the mixture of cells and enzymes can be directly added to prepared Concanavalin A beads, since solutions used in cell permeabilization for CUT&RUN contain EDTA, which inhibits the activity of collagenase and dispase enzymes.

**Figure 1.**
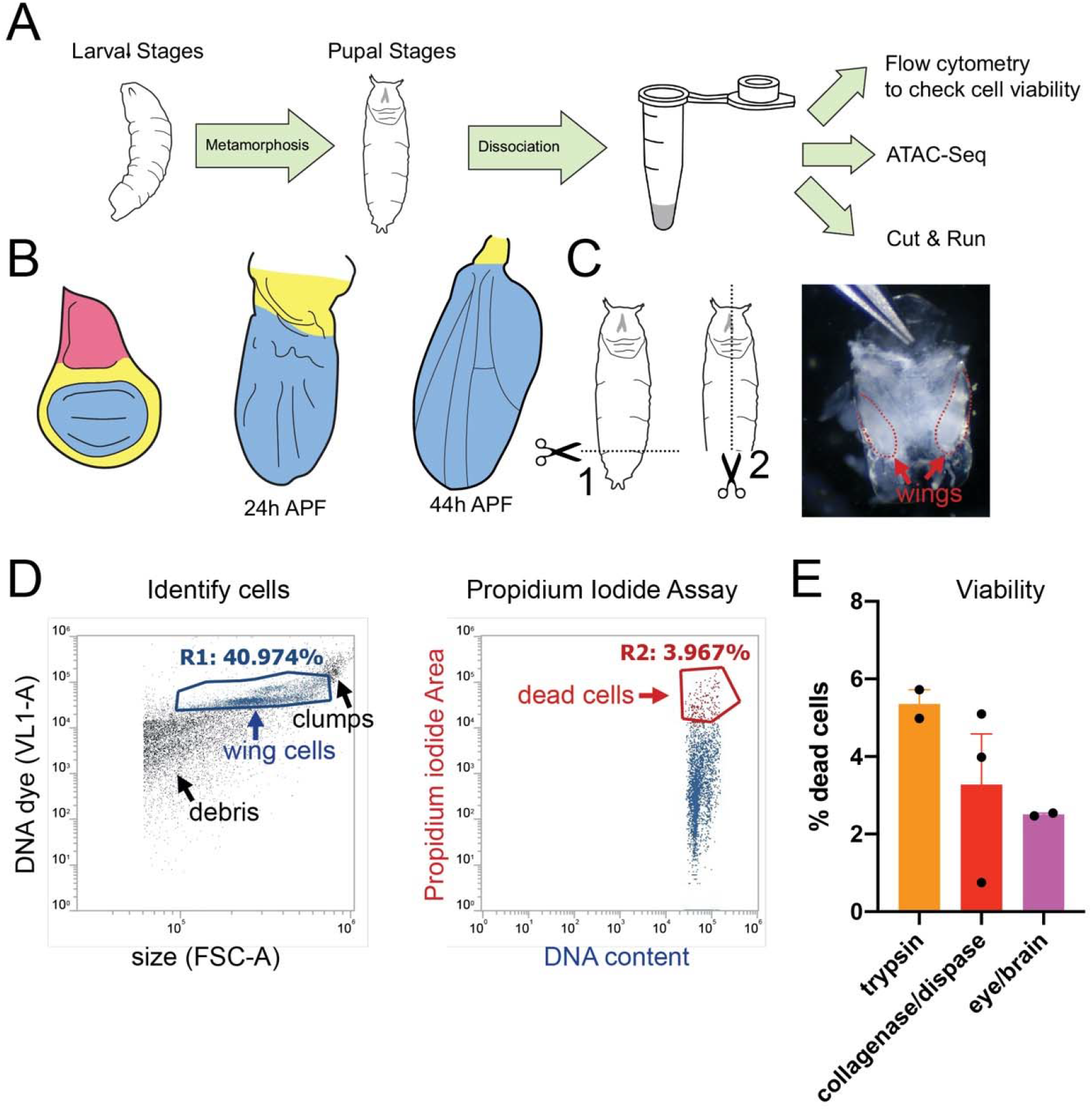
A wing dissociation protocol to release cells from cuticle and minimize cell death and loss. **(A)** Workflow diagram of assays compatible with gentle dissociation on larval and pupal tissues. Pupal tissues after 6h into metamorphosis are encased within pupa cuticle and require dissociation for the subsequent assays. **(B)** Diagrams of pupal wing morphogenesis during metamorphosis. Notum (pink) is present in larval wings but absent from dissected pupa wings after 18h APF. Larval and pupal wings contain hinge (yellow) and wing pouch (blue). **(C)** Diagram of pupa dissection (dotted lines) with image of 24h APF pupa removed from the tanned pupal case. Wings are enclosed in shiny, translucent pupa cuticle and are manually dissected from the body at the hinge for dissociation. **(D)** Example flow cytometry plot of dissociated 24h APF pupal wings. Cells were stained with a vital DNA dye (DyeCycleViolet) to discern cells from debris. Cell viability was assayed using a Propidium Iodide (PI) permeability assay and dead or dying cells were quantified based on gating of PI positive cells. **(E)** Quantifications of dead/dying cells in trypsin-based dissociation vs. collagenase/dispase dissociation in 24h APF pupal wings and 24h APF pupal eye/brain complexes.

To assess whether our tissue dissociation protocol is compatible with chromatin accessibility assays via ATAC-seq, we first compared ATAC-seq data we obtained from 3 replicates of 10 dissociated pupal wings across L3, 24h and 44h APF timepoints to our previously published FAIRE-seq data on 3 replicates of 40 wings across the same timepoints (Fig. 2). We see good agreement of the ATAC-seq data on dissociated wings with 58% of identified accessible peaks overlapping with peaks in our previous FAIRE-seq data on wings and most peaks exhibiting similar opening and closing dynamics over time. This demonstrates that after tissue dissociation, comparable chromatin accessibility profiles can be obtained with ¼ of the input used for FAIRE-seq. We also observe that ATAC-seq picks up more accessible peaks within gene coding regions, while FAIRE peaks are more likely to be at promoters, which was also previously observed in a direct comparison of FAIRE-seq and ATAC-seq signals in the *Drosophila* larval eye (Davie *et al.*, 2015).

**Figure 2.**
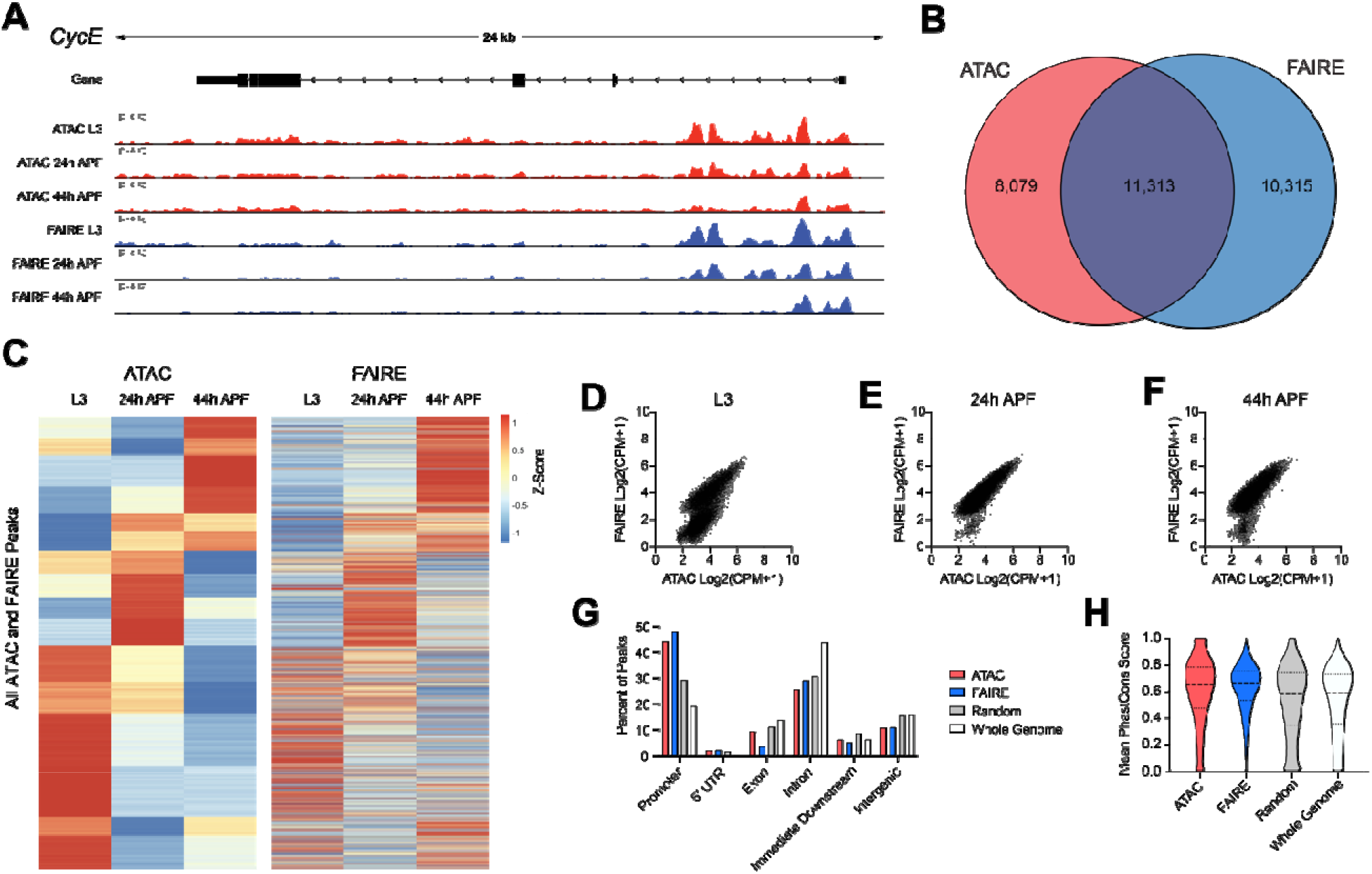
ATAC-Seq libraries from larval and pupal wings mirror accessibility profiles generated using FAIRE-Seq. (**A**) Accessibility profiles at the *cycE* gene locus generated using ATAC-Seq (red) and FAIRE-Seq (blue) from third larval instar (L3) wing discs and pupal wings at 24h and 44h after puparium formation (APF). Arrows on gene diagram indicate the direction of transcription. Y-axes for ATAC-Seq and FAIRE-Seq tracks: normalized read counts per million. (**B**) Venn diagram depicting the overlap between ATAC-Seq and FAIRE-Seq peaks called from any time point. (**C**) Heatmap depicting the average signal intensity dynamics at the union set of all peaks defined by ATAC-Seq and FAIRE-Seq. Data are scaled by Z-score and hierarchically clustered based on ATAC-Seq dynamics. (**D-F**) Scatter plots depicting the signal intensity (log2-transformed read counts per million) in FAIRE-Seq (y-axis) and ATAC-Seq (x-axis) libraries from L3 discs (D), 24h APF (E), and 44h APF (F) wings. Each plot includes th**e** union set of peak regions called at the given time point. (**G-H**) Percent of peaks residin**g** at various genomic elements (G) and distribution of mean PhastCons scores (H) from ATAC-Seq (red) or FAIRE-Seq (blue) libraries, as well as from randomized peak regions (grey) and 250 bp regions covering the entire genome (white). ATAC-Seq and FAIRE-Seq datasets include peaks called at any time point.

While tissue dissociation is essential for ATAC-seq on pupa wings after pupa cuticle formation (after 6h APF), ATAC-seq without tissue dissociation has been previously employed on larval wings, which are not yet enclosed within cuticle (Harris et al., 2020; Vizcaya-Molina et al., 2018). We therefore compared ATAC-seq profiles on dissociated larval wings with undissociated larval wings to assess how the dissociation process may impact chromatin accessibility assays (Fig 3). In general, we find good agreement between datasets from dissociated and undissociated tissues of the same stage. However, we noted ATAC-seq libraries prepared from dissociated wings required fewer PCR cycles of amplification, exhibited fewer reads from mitochondria and *Wolbachia* and better rates of reads mapping to the *Drosophila* genome, suggesting improved sensitivity of the assay after tissue dissociation. We suggest this may be due to improved access of the transposase enzyme to all nuclei of the tissue after dissociation, rather than just those on the surface of undissociated tissues. We therefore propose that dissociation may also assist with improving sensitivity and recovery from enzyme-based chromatin assays in thick tissues, in addition to allowing access to those with significant extracellular matrix or cuticle deposition. We have successfully performed this assay with dissociation on larval and pupal eyes and brains with similar sensitivity. Since tissue dissociation is essential for ATAC-seq on pupal tissues, we suggest that whenever comparisons between larval and pupal tissues are performed, larval tissues must also be dissociated.

**Figure 3.**
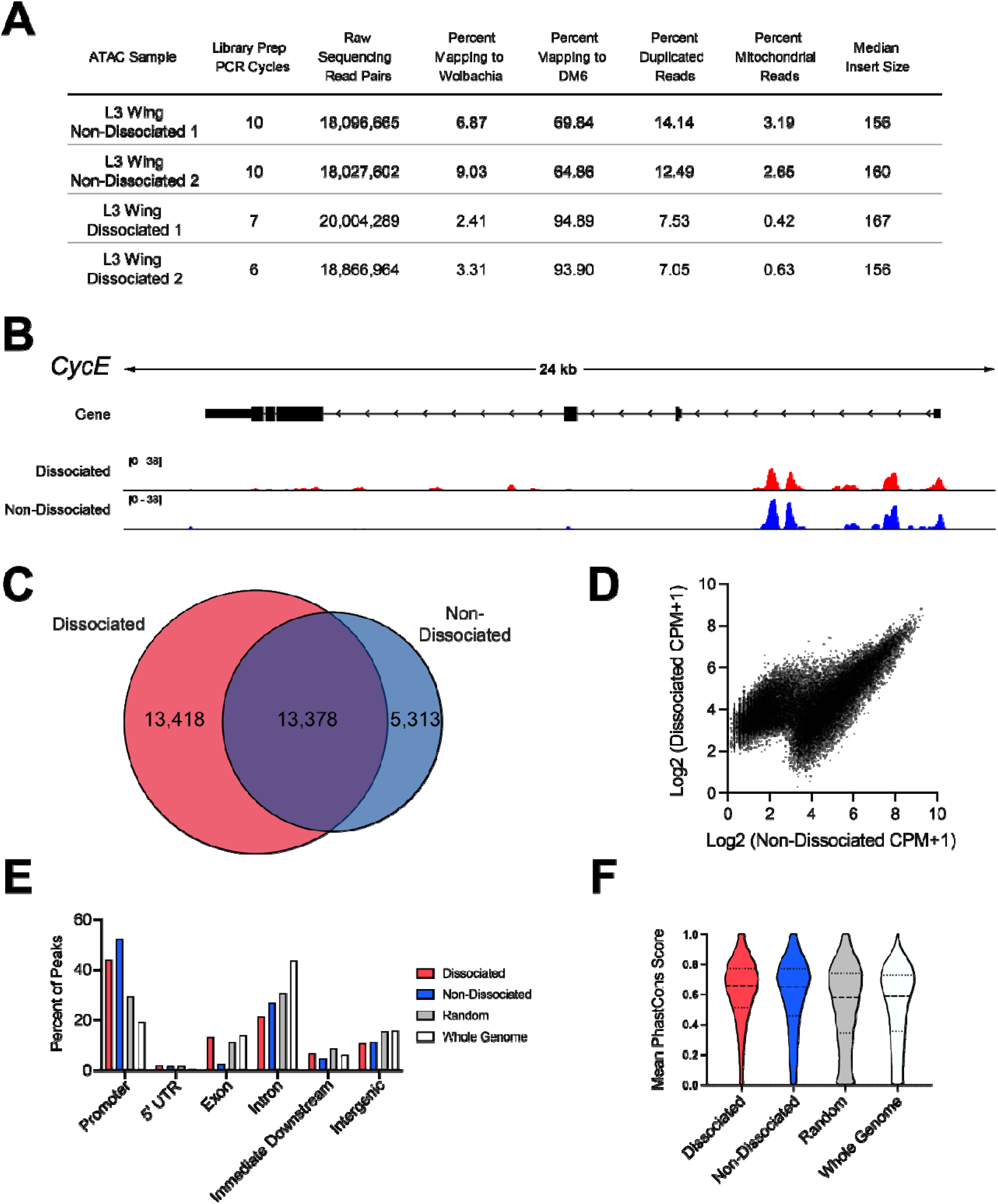
Wing disc dissociation is compatible with generation of high-quality ATAC-Seq libraries. (**A**) Table of quality metrics comparing ATAC-Seq libraries prepared from non-dissociated versus dissociated third larval instar (L3) wing discs. (**B)** ATAC-Seq reads at the *cycE* gene locus from dissociated (red) and non-dissociated (blue) L3 wing discs. Arrows on gene diagram indicate the direction of transcription. Y-axes for ATAC-Seq tracks: normalized read counts per million. (**C**) Venn diagram depicting the overlap between ATAC-Seq peaks called from dissociated (red) and nondissociated (blue) L3 wing discs. (**D**) Scatter plot depicting the average signal intensity (log2-transformed counts per million) in dissociated (y-axis) and non-dissociated (x-axis) L3 wing discs for the union set of peak regions called in each condition. (**E-F**) Percent of peaks residing at various genomic elements (E) and distribution of mean PhastCons scores (F) from dissociated (red) or non-dissociated (blue) L3 wing disc ATAC-Seq libraries, as well as from randomized peak regions (grey) and 250 bp regions covering the entire genome (white).

We next tested whether our dissociation protocol was compatible with the enzyme-based CUT&RUN assay to examine histone modification distribution in larval and pupal wings. To date, the only approach to examine the genome-wide localization of chromatin binding proteins or histone modification distribution in pupal wings after 6h APF has been Chromatin IP, which required a massive dissection of 700-1,000 pupa wings (Uyehara et al., 2017). We performed wing dissociation followed by CUT&RUN to localize the repressive histone mark Histone H3 Lysine 27 trimethylation (H3K27Me3) on L3 larval and 24h APF pupal wings with inputs ranging from 20-75 wings per sample (Fig. 4). Even with an input of only 20 wings (~1 million cells), we readily detect regions of broad H3K27Me3 that are consistent with known H3K27Me3 signals identified by ChIP in larval wings (Loubiere et al., 2016). Unfortunately, there are no published H3K27Me3 ChIP datasets from the *Drosophila* pupal wing available for comparison, but when we examine known temporally stable, wing-specific regions of H3K27Me3 enrichment (e.g. the Bithorax Complex), we observe the expected patterns of this histone mark across inputs ranging from 20-75 wings at both larval and pupal stages (Papp and Muller, 2006).

**Figure 4.**
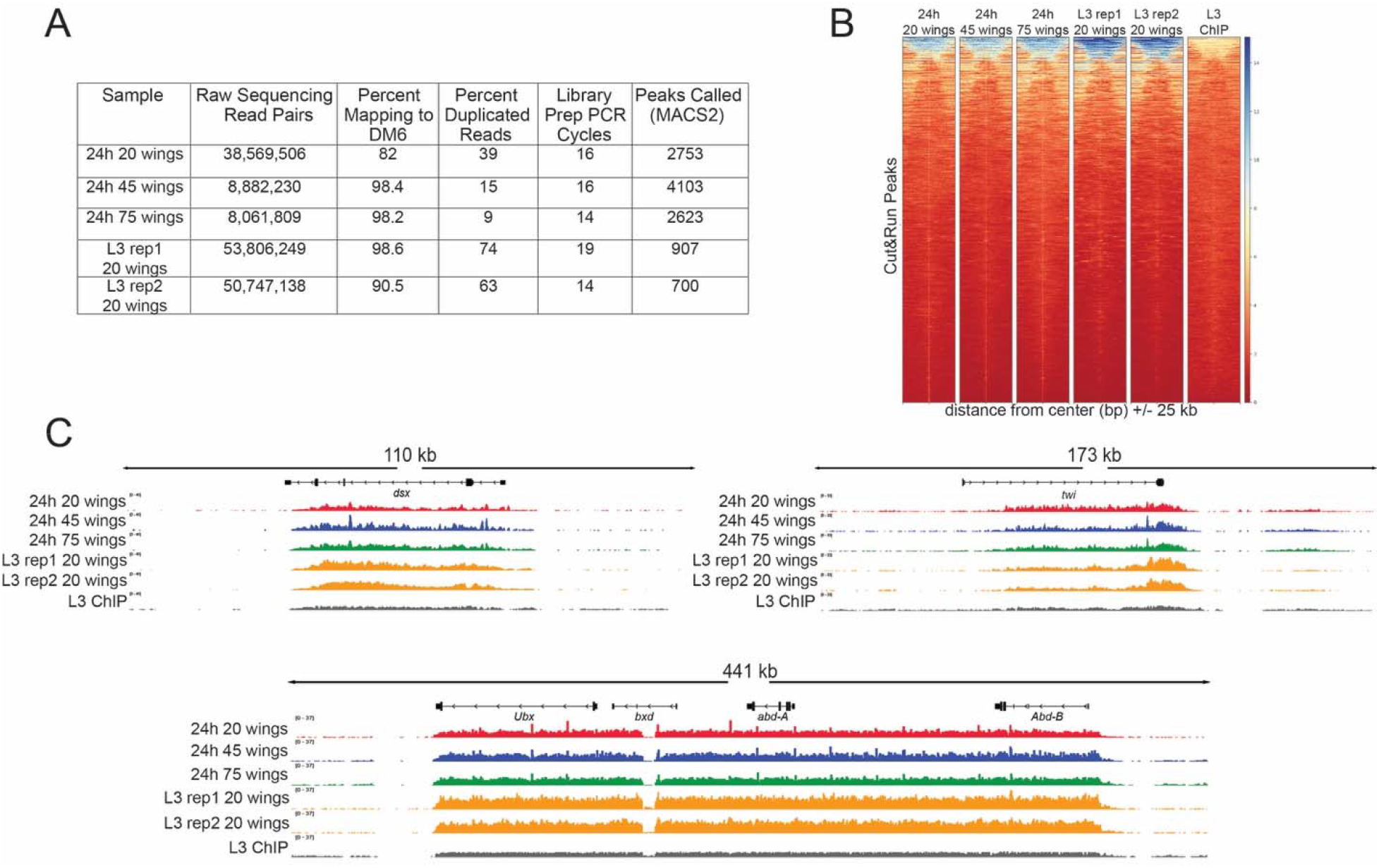
H3K27me3 CUT&RUN on dissociated pupal wings identifies stable domains similar to H3K27me3 ChIP-seq on larval wings. (**A**) Table of quality metrics comparing H3K27me3 CUT&RUN data prepared from 20 (red), 45 (blue) and 75 (green) dissociated 24h APF wings and third larval instar dissociated wings (gold) to H3K27me3 third larval instar wing ChIP-seq (black). Downsampling of the CUT&RUN 20 wings and third larval instar samples was performed to account for the varying sequencing depth; number of peaks shown reflects downsampled data. (**B**) Heatmaps showing signal intensity of peaks called from CUT&RUN dissociated 24h APF wings with differing sample input compared to peak signal intensity in third larval instar ChIP-seq. (**C**) Browser tracks showing H3K27me3 coverage at *dsx, twi,* and the bithorax complex containing *Ubx, bxd, abd-A* and *Abd-B* loci comparing CUT&RUN wing samples with third larval instar wing disc ChIP-seq. Browser tracks were group autoscaled.

## Summary

The tissue dissociation protocol described here is optimized for cell recovery and viability for use with subsequent ATAC-seq and CUT&RUN assays. This protocol will greatly facilitate genome-wide examinations of chromatin accessibility, histone modifications, and localization of chromatin binding proteins in tissues across metamorphosis at timepoints previously inaccessible to these assays.

## Acknowledgements

Work in the Buttitta lab is supported by the NIH/NIGMS (R01) GM127367. Work in the McKay Lab is supported by NIH/NIGMS (R35) GM128851. EMB was supported by the U.Michigan Genetics Training Grant (T32GM007544). We thank Matt Niederhuber for kindly providing the illustration in Fig. 1 adapted from (Niederhuber and McKay, 2021). We thank Dr. Steven Henikoff (FHCRC) for providing pA/G MNase. We thank Dr. Yiqin Ma for initial studies into ATAC-seq on pupal wings.

